# Old Goats: 3,000 years of genetic connectivity of the domestic goat in Ireland

**DOI:** 10.1101/2025.09.26.678852

**Authors:** Judith Findlater, Jolijn Erven, Alex Siekmann, Valeria Mattiangeli, The Vargoats Consortium, Eileen Murphy, Kevin G. Daly

## Abstract

The domestic goat likely first arrived to the island of Ireland as part of the introduction of agriculture approximately 5,900 years ago, and remains a part of the island’s biocultural heritage. However, due to the challenges of differentiating goat remains from that of sheep using traditional archaeozoological approaches, there are few specimens specifically identified as goat. To address this we employed genetic, proteomics, and archaeozoological techniques to assess faunal remains from the Late Bronze assemblage of Haughey’s Fort (Armagh) and medieval assemblage of Carrickfergus (Antrim). We identify these specimens as goats using proteomics and genetics, and additionally determine their molecular sex.

Genomic data recovered from a Haughey’s Fort goat reveals a three-millenia genetic connection between herds in the Late Bronze Age, medieval period, and the indigenous Irish breed extant today, the Old Irish Goat. We additionally find varying levels of inbreeding within goats from the settlement of Carrickfergus, suggesting possible mixed use of herds within medieval Irish society. Our results demonstrate the continuing potential of combining archaeological and biomolecular techniques to clarify existing ambiguities and at the same time reveal new facets of the past.

## 1. Introduction

Livestock management on the island of Ireland began 5,900-5,750 years ago with the arrival of agricultural practices and people (Shennan, 2018; Sheridan, 2010; Whitehouse et al., 2014). Among the introduced species was, assumedly, the domestic goat *Capra hircus* (Porter et al., 2016), which today retains a place in Ireland’s traditions, placenames, art, and biocultural heritage (Mac Coitir, 2010; Mac Con Iomaire, 2014). While there are several potential goat remains in Ireland’s earliest domestic faunal assemblages in the Neolithic period, they are typically undifferentiated from sheep; the enigmatic “ovicaprid”, “caprovine” or “sheep/goat” is a persistent challenge in zooarchaeological research (Seabrook et al., 2025). Examples of some ambiguous assignments include remains from Audleystown, Ashleypark, Ballyalton, Fourknocks, and Tankardstown (McCormick, 1985). Thus, there are no unambiguously identified goats in the Irish Neolithic archaeozoological record and none are identified in the Beaker/Early Bronze Age period (McCormick, 2007). The Late Bronze Ages assemblages of Haughey’s Fort (Armagh) and Mooghaun (Clare) are one of the first to offer definitive evidence for the presence of goats ∼1,000 cal BCE (McCormick, 2007). Other hints of the presence of goats come from genetic data recovered from bog butter from Early Bronze Age Knockdrin (1740-1715 cal. BCE, (Mattiangeli et al., 2020)).

The difficulty in differentiating sheep and goat postcranial remains based on archaeozoological methodologies has long been recognized (Boessneck, 1969; Cornevin and Lesbre, 1891; Payne, 1969). A substantial body of research has been undertaken to identify diagnostic features in caprine dentition, although these have had mixed success (Gillis et al., 2011; Zeder and Lapham, 2010). Recent decades have seen improvements in the tools available to tackle this challenge; for example, novel measurement approaches in combination with morphological approaches (Salvagno and Albarella, 2017). Petrousal parts of the temporal bone can be assigned to either species, despite intraspecific variation (Mallet et al., 2019). Geometric morphometrics may also provide distinguishing power e.g. (Haruda, 2017; Jeanjean et al., 2022; Lloveras et al., 2022). Molecular methods such as type I collagen protein sequencing (Buckley et al., 2010) or ZooMS (Buckley et al., 2009; Seabrook et al., 2025) additionally provide a powerful means of positively identifying either sheep or goat from small amounts of archaeological materials.

A previous study has identified a substantial degree of matrilineal connection between historical specimens of the “Old” landrace breeds of England, Ireland and Scotland (Cassidy et al., 2017). Among mitochondrial genomes of “Old Irish Goats” today, their sequences group either with this historical clade, or among the broader diversity of European goats. The question remains as to the age of this “British and Irish” historical grouping - if it represents the long-standing herds of the islands, likely introduced in the Neolithic period approximately 6,000 years ago (Yalden, 1999), or more recent demographic events or movements of animals by humans.

This paper investigates the remains of eight goats obtained from the Late Bronze Age site of Haughey’s Fort (Armagh) and the medieval town of Carrickfergus (Antrim) identified using archaeological approaches, to determine if molecular approaches applied to the archaeological record can help resolve this issue. Using a combination of proteomic and genomic approaches, we identify domestic goats from among the eight samples, and assess their relationship to goat breeds today. The animals from Haughey’s Fort are the oldest verified goats to have been identified in Ireland to date.

## 2. Materials and Methods

### 2.1 Faunal remains

The faunal remains included in the study derive from the Late Bronze Age site of (Armagh) and the medieval urban settlement of Carrickfergus (Antrim). Ovicaprid skeletal elements from both sites were subjected to morphological (Boessneck, 1969; Payne, 1985; Prendergast et al., 2019; Prummel and Frisch, 1986; Zedda et al., 2017; Zeder and Lapham, 2010) and metrical (Davis, 2017; Gron et al., 2019; Rowley-Conwy, 1998; Salvagno and Albarella, 2017) analysis in Queen’s University Belfast for the purpose of identifying goat bones with the intention that this could then be confirmed using ZooMS.

Haughey’s Fort dates to 1,100-900 BCE and it lies within the Navan complex of sites, some 1 km to the west of the famous royal site of Navan Fort. It is a trivallate hill fort with an inner ditch enclosing an area with a breadth of approximately 150 m. The inner ditch (Tr 5) was waterlogged and it produced a wealth of organic remains, including a substantial collection of animal bones (Mallory et al., 1996). The assemblage was dominated by cattle (MNI=69), with pigs of secondary importance (MNI=43). Sheep/goat were present in very small numbers with an MNI of 4. Of particular note was the presence of very long horned goats, with two near complete horn cores having lengths of 344 mm and 348 mm (with the latter broken at its tip). The presence of a small number of large cattle and dogs was interpreted as indicating that selective breeding of large animals had occurred (Murphy and McCormick, 1996). The three elements selected from Haughey’s Fort for inclusion in the study all derived from the waterlogged inner ditch. One of the large horn-cores (HF89, IC/21) was also included but, since its morphology was so definitively goat, it was not considered necessary to subject it to ZooMS analysis.

The town of Carrickfergus overlooks Belfast Lough and is situated on the southern coast of County Antrim. It has been an urban settlement for the past 800 years since its foundation by the Anglo-Norman, John de Courcy, in the late 1170s CE. It has been the subject of numerous excavations since the 1970s (Ó Baoill, 2008). Three of the ovicaprid bones from Carrickfergus derived from excavations undertaken in Antrim Street (CFXIV) in advance of development in the 1980s. Midden-like layers identified as 13th-16th century in date containing a substantial quantity of animal bone, and a possible animal feeding trough located within a lean-to structure, were discovered (Brannon, 1980-1984). The CFXIV assemblage was dominated by cattle (MNI=49), with ovicaprids of secondary importance (MNI=33) and pigs also well represented (MNI=18) (Findlater, 2021). The fourth sample was obtained from excavations undertaken in 1991 in advance of development in an enclosed courtyard building to the rear of 25 West Street (CF19). A large quantity of animal bone was recovered and the ovicaprid included in the study derived from a pit (F49) dated to the 13-14th centuries (Ó Baoill, 1993). The assemblage from this era was dominated by cattle (MNI=25), with ovicaprids (MNI=9), dog (MNI=6) and pig (MNI=5) also reasonably well represented (Murphy, 2012).

### 2.2 ZooMS

Seven of the eight samples were sent to BioArCh, University of York, for Zooarchaeology by Mass Spectrometry (ZooMS) analysis. Fragments were taken by drilling a small part of each element. The sample site where the drilling occurred was selected to cause minimal damage to the skeletal element. The samples were submitted for ZooMS analysis where they were prepared in the laboratory to extract peptides from collagen samples (Buckley et al., 2010, 2009; Hendy, 2021). Using a MALDI-Time-of-Flight (TOF) mass spectrometer, the ‘fingerprint’ of various peptides was analysed and through the comparison of the peptide masses generated, species identification was determined for each of the samples submitted (Hendy, 2021). It is possible to differentiate between sheep and goats based on a single peptide with two amino acid differences. In the Time-of-Flight spectra the goat marker peptide occurs at 3093 *m/z*, while the sheep peptide is evident at 3033 *m/z* (Table S2 ZooMs, Figure S1 (Buckley et al., 2010)).

### 2.3 Radiocarbon Dating

Two specimens were subject to radiocarbon dating, Carrick3/IRCA121 and Haughey1/IRHA201. Dating was performed at the ^14^Chrono Centre (Belfast) and Oxford Radiocarbon Accelerator Unit (Oxford) for Haughey1 (approximately 1000mg) and Carrick3 (137mg) respectively. Oxcal (Bronk Ramsey, 1994) and INTCAL20 (Reimer et al., 2020) were used to calibrate ages.

### 2.4 DNA extraction

All eight bone and tooth remains were sampled for DNA extraction in dedicated ancient DNA laboratory facilities in Trinity College Dublin. Approximately 100mg was obtained using a dental blade for each sample, cutting either the densest part of the bone element or from the available roots of each tooth. Specimens were then powderized using a MixerMill (Retsch).

DNA extraction followed a previously described pipeline (Mattiangeli et al., 2023),. Briefly, samples were subject to a dilute sodium hypochlorite (0.5%) wash, followed by three H_2_O washes, and short EDTA predigestion (Damgaard et al., 2015) followed by an EDTA-proteinase K overnight digestion. The resulting supernatant was purified using High Pure Viral Nucleic Acid Large Volume purification kits Roche), eluting DNA in 50μL of EBT (7.5μL of 20% Tween in 15mL EB buffer).

Purified DNA was then used to construct double stranded DNA sequencing libraries (Meyer and Kircher, 2010). Uracil bases were first excised (Briggs et al., 2010) by treating DNA with 5μL Uracil-DNA glycosylase (UDG) over 3 hours at 37°C. Sample IRHA201 was erroneously not subject to dry-spin step during extraction, so 26.25 µL extract was incubated with 8 µL UDG enzyme instead. DNA library construction then followed steps outlined in (Daly et al., 2018). Amplification (12 cycles) was performed using Accuprime DNA polymerase (Invitrogen), incorporating dual sequencing oligo indexes. Amplified DNA was quantified using a Tapestation 4000 (Agilent). Additional DNA amplifications were performed at amplification cycles matching an expected 1ng/μL concentration. The resulting amplifications were subject to high throughput sequencing on Illumina Novaseq 4000 (TrinSeq, Dublin) and NovaSeq X (Macrogen Europe, Amsterdam).

### 2.5 Data handling and genotyping

Adaptor trimming and read merging was performed using AdaptorRemoval version 2.3.2 (Schubert et al., 2016), removing collapsed reads shorter than 30bp and removing low quality terminal bases (--minadapteroverlap 1 --adapter1 AGATCGGAAGAGCACACGTCTGAACTCCAGTCAC --adapter2 AGATCGGAAGAGCGTCGTGTAGGGAAAGAGTGT --minlength 30 --trimns --trimqualities). Collapsed and raw read qualities were assessed using FastQC (Andrews and Others, 2010). Specimen species were determined using FastQ Screen (Wingett and Andrews, 2018), aligning reads against a set of genomes including sheep, goat, cattle, and human. We calculated coverage with Qualimap2 (Okonechnikov et al., 2016).

Following species ID (Table S1), reads from UDG-treated libraries of goat samples were aligned to the goat reference genome ARS1.0 (Bickhart et al., 2017) and using a pipeline previously described (Daly et al., 2021). Briefly, collapsed reads were aligned using bwa aln version 0.7.17-r1188 (Li and Durbin, 2009) relaxing alignment parameters (-n 0.01 -o 2) and converted to bam files using samtools version 1.13 (Li et al., 2009). Using samtools, bam files were filtered to remove reads with mapping quality <30, and for duplicate reads. Mismatches were calculated on the mitochondrial DNA (Table S3).

Pseudohaploid genotypes for variant sites discovered in the VarGoats genome dataset (Denoyelle et al., 2021; Erven et al., 2025) were called by random read sampling using ANGSD (Korneliussen et al., 2014), previously described (Erven et al., 2025). Additionally, samples were imputed using GLIMPSE2 (Rubinacci et al., 2023) using a pipeline previously described (Erven et al., 2025), employing the VarGoats dataset as a reference panel and a recent 10kb-scale recombination map as the genetic map (Etourneau et al., 2025). Imputed samples were filtered for a genotype probability (GP) of 0.99.

### 2.6 LASER

A projection Principal Component Analysis (PCA) using LASER v2 (Wang et al., 2015) was performed to maximize the inclusion of low-coverage samples (e.g., Haughey1). The PCA reference space and projection transformation were constructed using the VarGoats dataset, filtered to retain transversions with a MAF of 5%, resulting in a total of 2,420,382 SNPs. All ancient samples were then projected onto the PCA space, and then filtered for individuals covered by less than 2000 loci. To reduce stochastic variation and provide robust estimates, 100 replicates were conducted.

### 2.7 Outgroup *f*_*3*_

To estimate the degree of shared genetic drift between ancient Irish goats and modern breeds, outgroup *f*_*3*_ statistics (Patterson et al., 2012) were calculated using ADMIXTOOLS version 7.0.2 (Maier et al., 2023) between medieval Ireland (Carrick1, Carrick3), Bronze Age Ireland (Haughey1), Bronze Age England (Potterne1), and modern Old Irish Goat and each domestic breed grouping in the VarGoats dataset. We used granular breed groupings where possible and domestic sheep were used as the outgroup. Outgroup *f*_*3*_ statistics were also calculated to estimate the degree of shared genetic drift between ancient Irish goats and other ancient populations. Ancient goats were grouped based on settlement and time period, Ganj Dareh was divided into a Ganj Dareh and Ganj Dareh wild group, due to a higher amount of wild ancestry in some of the Ganj Dareh goats (Daly et al., 2021).

### 2.8 *D* statistics

To test for evidence of gene flow and estimate its extent between Irish goats and modern breeds, *D* statistics were calculated using ADMIXTOOLS version 7.0.2 (Maier et al., 2023). The model to test for genetic differentiation between the modern Old Irish Goat breeds and ancient Irish/British goats was structured as *D*(Sheep, Modern breed, Old Irish Goat, ancient goats). To test for shared ancestry among ancient goats *D* statistics were calculated as *D*(Sheep, ancient Irish/British goat, ancient goats/modern Old Irish Goats, ancient goats/modern Old Irish Goats). The Blagotin goats were grouped and considered as one population.

### 2.9 Runs-of-homozygosity (ROH)

ROH were computed on imputed genotypes using a combined plink and bcftools pipeline, previously described and validated (Erven et al., 2025). In summary, samples were downsampled to 1.2 million SNPs, filtered for minor allele frequency >5% and restricted to transversions. ROH were calculated for each individual using both PLINK v1.90 (Chang et al., 2015) and bcftools v1.17 (Danecek et al., 2021). Plink was run with standard parameters aside from setting the --homozyg-window-snp to 200. Bcftools was run with standard parameters aside from -G 30 --AF-dflt 0.4. Bcftools output was further filtered to retain regions containing at least 200 SNPs, a quality score greater than 10, and a minimum length of 500kb. Long ROHs identified by bcftools (>4Mb) were merged with the plink ROH profiles using mergeBed (Quinlan and Hall, 2010), counting the number and keeping the sizes of merged ROHs with the parameters -c and -o count, collapse.

### 2.10 Identity-by-Descent (IBD)

Identity-by-descent segments were identified using a similar pipeline as described in *Erven et al*., *2025* (Erven et al., 2025). In summary, imputed genotypes were phased together using Beagle5 (Browning and Browning, 2013) with standard parameters aside from setting impute=true and specifying a random seed. Genotypes below GP99 were set to missing, and SNPs were further filtered for the 1,037,536 SNPs used in *Erven et al*., *2025* (MAF 5%, transversions and had no missing data in imputed samples >2×), and finally filtered for no missingness.

RefinedIBD (Browning and Browning, 2013) was run with default settings. To correct for breaks or short gaps in IBD segments, segments were merged with a sex-averaged 100kb-scale recombination map (Etourneau et al., 2025), using merge-ibd-segments.17Jan20.102.jar, allowing a maximum of one discordant homozygote and gaps less than 0.6 cM in length. The phasing, refinedIBD and merge-ibd were repeated a total of 3 times to account for potential variance in phasing. The first 2 Mb of chromosome 18 was excluded as in *Erven et al*., *2025*. The repeated runs were combined and filtered for a minimum LOD score of 3 and a length threshold of 3cM. To obtain IBD segments for a pair of individuals, IBD segments for pairs of individuals were extracted from the IBD files and merged with bedtools MergeBed.

### 2.11 Mitochondrial Enrichment

Two specimens (Carrick2 and Carrick4; Table S4) were subject to Mybaits in-solution RNA bait enrichment (Daicel Arbor Bioscience), using a previously reported capture array targeting domesticated species mtDNA (O’Sullivan et al., 2016). Amplified UDG-treated libraries were pooled for roughly equal representation, the pool was desiccated and then re-suspended in 8.4 μL H_2_O. RNA baits and blocks were added to the pool as manufacturer’s instructions, with one modification: Block #1 was replaced with 2.5μl of pooled DNA (total 8.4μL). Baits and DNA were incubated for 40 hours, 65°C. Captured DNA was recovered using Dynabeads® MyOne™Strepavidin C1 magnetic beads (ThermoFisher Scientific), resuspending in 30μL H_2_O. 15μL of the captured DNA was amplified (14 cycles) using KAPA HiFi DNA Polymerase (Kapa Biosystems), following the myBaits protocol. Captured libraries were sequenced on a NovaSeq 6000 (TrinSeq, Dublin).

### 2.12 Mitochondrial Genomes

Reads were aligned to a circularised version (Daly et al., 2018) of the goat mtDNA reference genome (NC_005044.2; 15bp of each side concatenated to the opposite end) using parameters described under “Data Handling and Genotyping”. mtDNA sequences were called using ANGSD version 0.937-74-g9100f3d (Korneliussen et al., 2014), using -doFasta 2 with the following parameters: -minQ 20 -minMapQ 30 -trim 4 -setMinDepth 3. For samples with <4× mtDNA coverage, -setMinDepth 1 was used instead. The resulting fasta files were decircularized using a python script by removing the first and last 15bp of the sequence. mtDNA sequences were aligned with published mtDNA sequences using MUSCLE (Edgar, 2004) and ML phylogenies constructed using phyML (Guindon et al., 2010) through Seaview (Gouy et al., 2010). After initial phylogenetic placement, samples were realigned to a closer haplogroup sequence (Daly et al., 2018) and final mtDNA sequences generated as above. A multiple sequence alignment was performed on the realigned mtDNA sequences with MAFFT (Katoh et al., 2002) and ML phylogenies were constructed with RAxML (Stamatakis, 2014), with 1000 bootstrap replicates. Modern mitochondrial sequences were retrieved from (Colli et al., 2015), ancient and historical mitochondrial sequences were retrieved from (Cassidy et al., 2017; Daly et al., 2018).

### 2.13 Molecular Sex

Molecular sex was determined using the relationship between the number of aligning reads to each chromosome vs. chromosome length (Park et al., 2015). Male samples compared to female samples are expected to show lower read alignment to the X chromosome relative to its length, due to the single copy of the X chromosome expected to be carried.

## 3. Results

### 3.1 Species identification

All seven specimens subject to ZooMS were identified as domestic goat *C. hircus* (Table S1, Figure S1). DNA sequencing data provided consistent species identification of six of these seven; the seventh sample did not have preserved endogenous DNA for genetic species ID (Haughey2). Additionally, genetic data provided species assignment for one sample not subject to ZooMS (Haughey4), confirming its archaeozoological identification as goat. Sequencing data displayed characteristics of ancient DNA, showing residual post-mortem damage after Uracil DNA-glycosylase treatment (Figure S2) and short read lengths (Figure S3, Table S1), albeit with a single specimen (Carrick1) showing longer read lengths (average 105 bp, compared to 56-75 bp of other Carrickfergus samples, Table S1). Molecular sex could be determined for all specimens with species ID; notably, all goats from Carrickfergus were female and those from Haughey’s Fort, male (Figure 1; Table S1).

**Figure 1.**
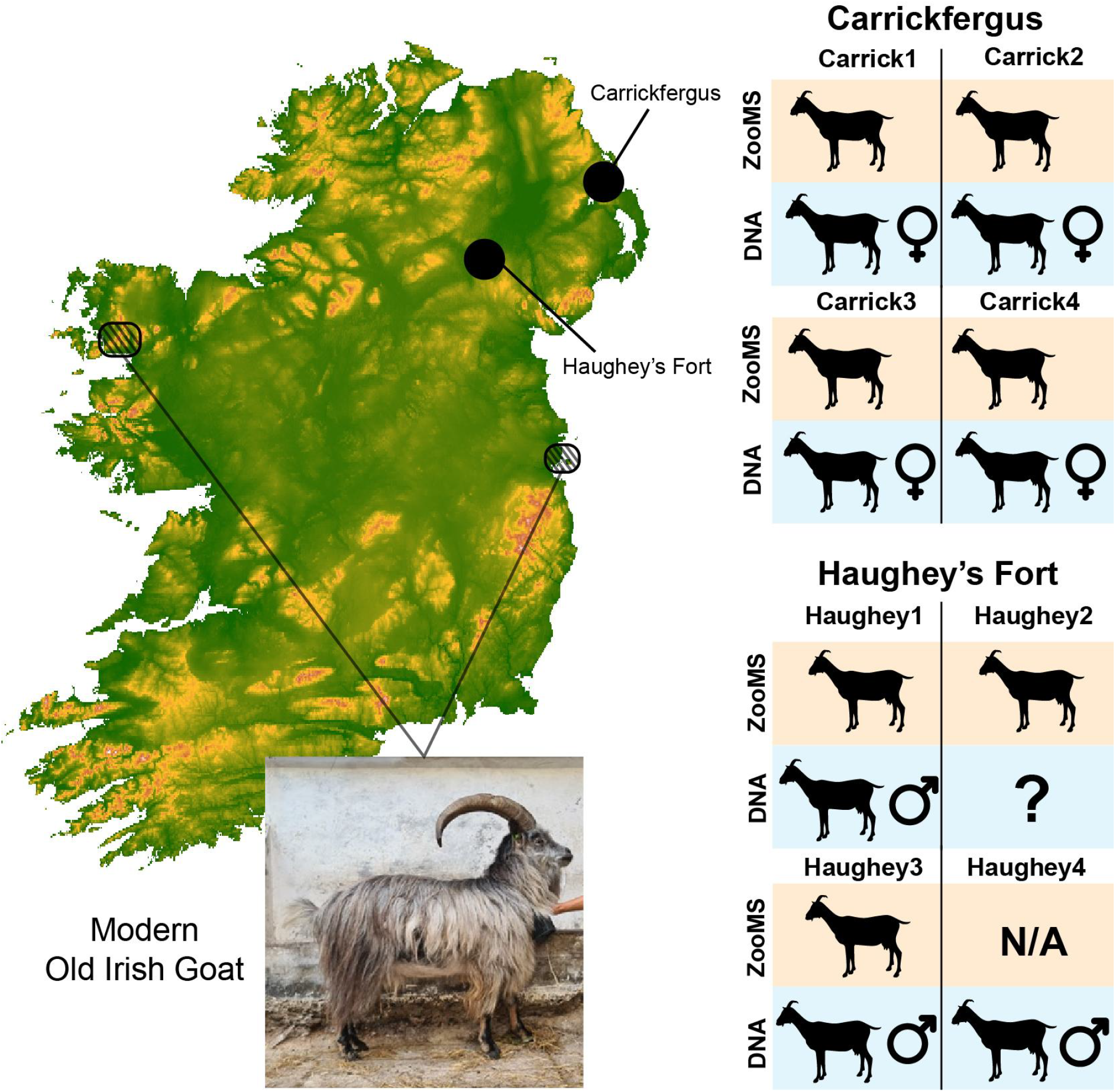
Summary of study area, species identifications and molecular sex results. A) map of the island of Ireland with the studied assemblages indicated (Haughey’s Fort and Carrickfergus), in addition to the regions in which the Old Irish Goat are primarily found today (Mulranny and Howth). Genetic and proteomic assessment of studied specimens. ? indicates samples for which data was not sufficient for molecular sex determination. Photo credit: The Old Irish Goat Society.

### 3.2 Radiocarbon Dating

Calibration of radiocarbon dates obtained for Haughey1 and Carrick3 (Figure S4; Table S1) confirmed the expected age of each. Haughey1, deriving from a Late Bronze Age context, dates to 1,112-947 cal BCE (UBA-55267; 2σ range). Carrick3, from the medieval settlement of Carrickfergus, dates to roughly the 16th century CE (OxA-44984; 1477-1635 cal CE), predominantly overlapping with the expected stratigraphy (13th-16th century CE).

### 3.3 Population genetic analyses

Sufficient sequencing data (Table S1) was available for population genetic analyses of three goat specimens: one Late Bronze Age goat from Haughey’s Fort (Haughey1) and two Late medieval goats from Carrickfergus (Carrick1 and Carrick3). Projection using LASER (Wang et al., 2015) placed these specimens with the general European section of the PCA (Figure 2). Specifically, the historic goats from Carrickfergus fell close to both modern Old Irish Goat genomes and a published Bronze Age goat (Potterne1) from Wiltshire, Britain (Daly et al., 2018). Haughey1 also falls within the general European section of the PCA; however the limited number of covered sites likely introduced bias in their placement, which we present with standard deviation to demonstrate the large uncertainty between LASER replicates.

**Figure 2.**
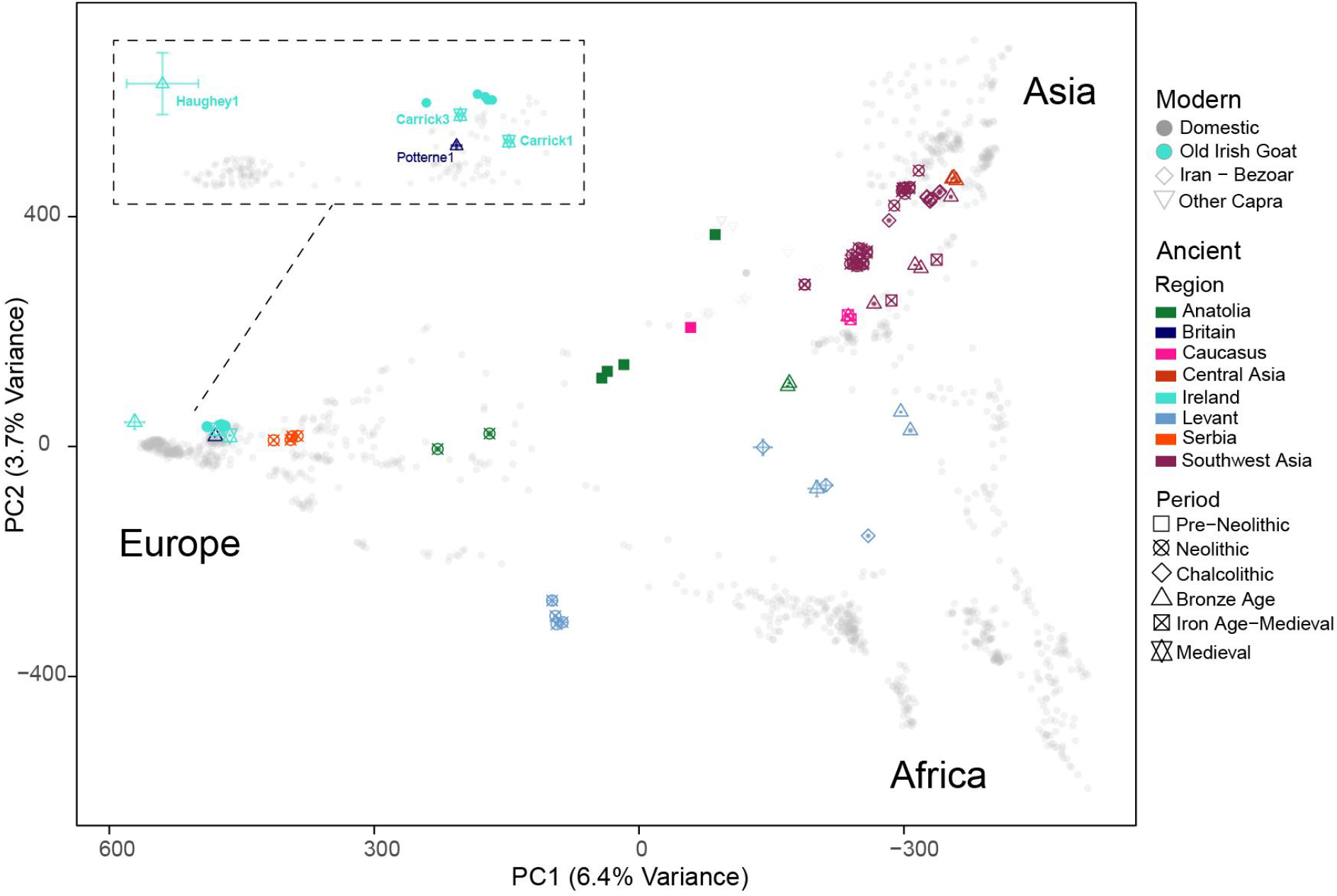
Projection PCA using Laser (Wang et al., 2015), placing ancient Irish goat genetic data in the context of modern and other ancient genomes. Subpanel provides a zoomed-in view of the PCA space in which the ancient Irish goat samples fall.

To assess the relationship of these pre-modern goats with goats from Ireland today, we computed measures of shared genetic drift (outgroup *f*_*3*_, (Patterson et al., 2012)) of the ancient genomes with each modern breed grouping in the VarGoats dataset (Table S5). For all three ancient goats, we found the highest shared drift with Old Irish Goat (Figure 3, Figure S5 for Carrick3), albeit with high uncertainty for the Haughey Fort individual (Table S6), a consequence of its paucity of aligned sequencing reads. However, outgroup *f*_*3*_ for goats of European origin are robust to low coverages (Supplementary note), supporting a genuine shared genetic history between Old Irish Goat and the Late Bronze Age genome.

**Figure 3.**
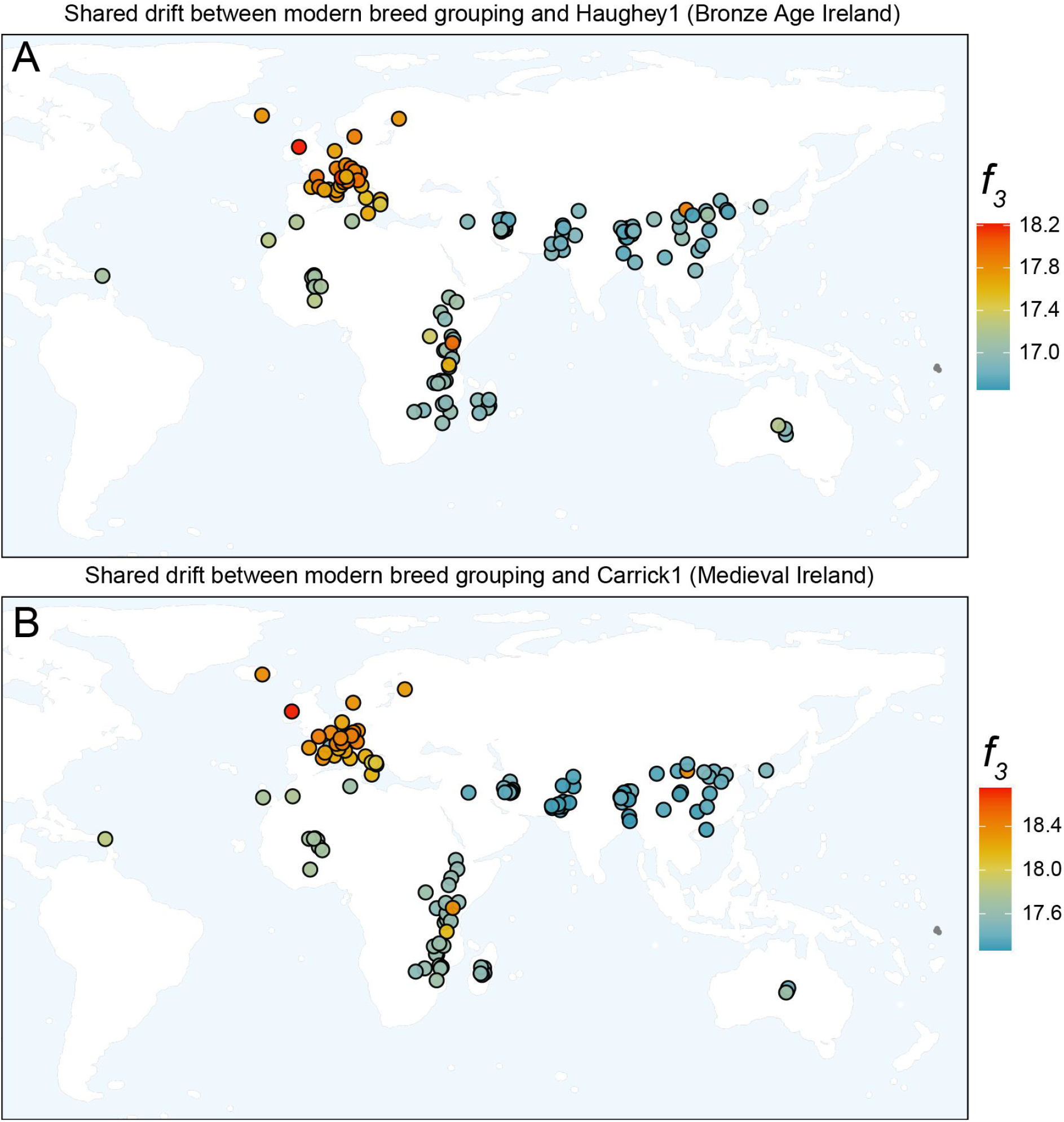
Geographical distribution of shared genetic drift (outgroup *f*_*3*_) between ancient Irish goats and modern breeds today. A) Shared genetic drift between a Bronze Age goat from Haughey’s Fort, Ireland (Haughey1), and modern domestic goat breeds. B) Shared genetic drift between a medieval goat from Carrickfergus, Ireland (Carrick1), and modern domestic goat breeds.

We next assessed shared genetic drift between ancient goats (Figure S6-8; Table S7), Haughey1 was excluded in the Carrick comparisons due to low number of reads. The Late medieval Carrickfergus goats have high pairwise affinities with each other and a Bronze Age British goat from Wiltshire (Potterne1, Britain), followed by other ancient European goats. We also assessed ancient goat affinity with the Late Bronze Age Haughey1 goat, and found its highest pairwise affinities with the Carrick3 individual, then Carrick1 and Potterne1, supporting temporal connectivity between Late Bronze Age and Late medieval Irish goat herds. We note deflated pairwise outgroup *f*_*3*_ values involving Carrick1 when calculated using pseudohaploid genotypes on transversions (Figure S6-8), which is mitigated using a minor allele frequency threshold of 5% (Figure S9-11), indicating that alleles with low frequency in the dataset introduced noise in our *f*_*3*_ estimates. *D* statistics mirror this trend, where the Carrickfergus goats show a high genetic affinity with each other, the British Bronze Age Potterne1 goat, and the modern Old Irish Goat breed (Table S8). This pattern is consistent across both Carrick1 and Carrick3, with Carrick3 showing a slightly higher affinity with Potterne1 and the Old Irish Goat breed, in line with the outgroup *f*_*3*_ results.

### 3.4 Mitochondrial phylogeny

The mitochondria of the Late Bronze Age Irish goat (Haughey1) and Late medieval Irish goats (Carrick1-4) fall within the global European goat diversity, primarily represented by the ubiquitous haplogroup A (Figure 4, S12 (Colli et al., 2015; Naderi et al., 2007)). However, none of the ancient Irish and Britain samples cluster within the two previously identified clades of historical Irish and Scottish samples (Cassidy et al., 2017). Haughey1 groups with a Swiss breed — albeit with a deep node, possibly due to the limited data available for this sample — while Carrick4 clusters with both a historical (H-Irish2) and a modern Irish goat (M-Irish4). Carrick3 falls on the outside of this cluster grouping with a Swiss Grisons Striped goat. The Carrick1 mitochondria groups with a Modern Irish goat (M-Irish3) and Carrick2 falls within a more diverse cluster of Swiss, Spanish and Austrian modern goat breeds.

**Figure 4.**
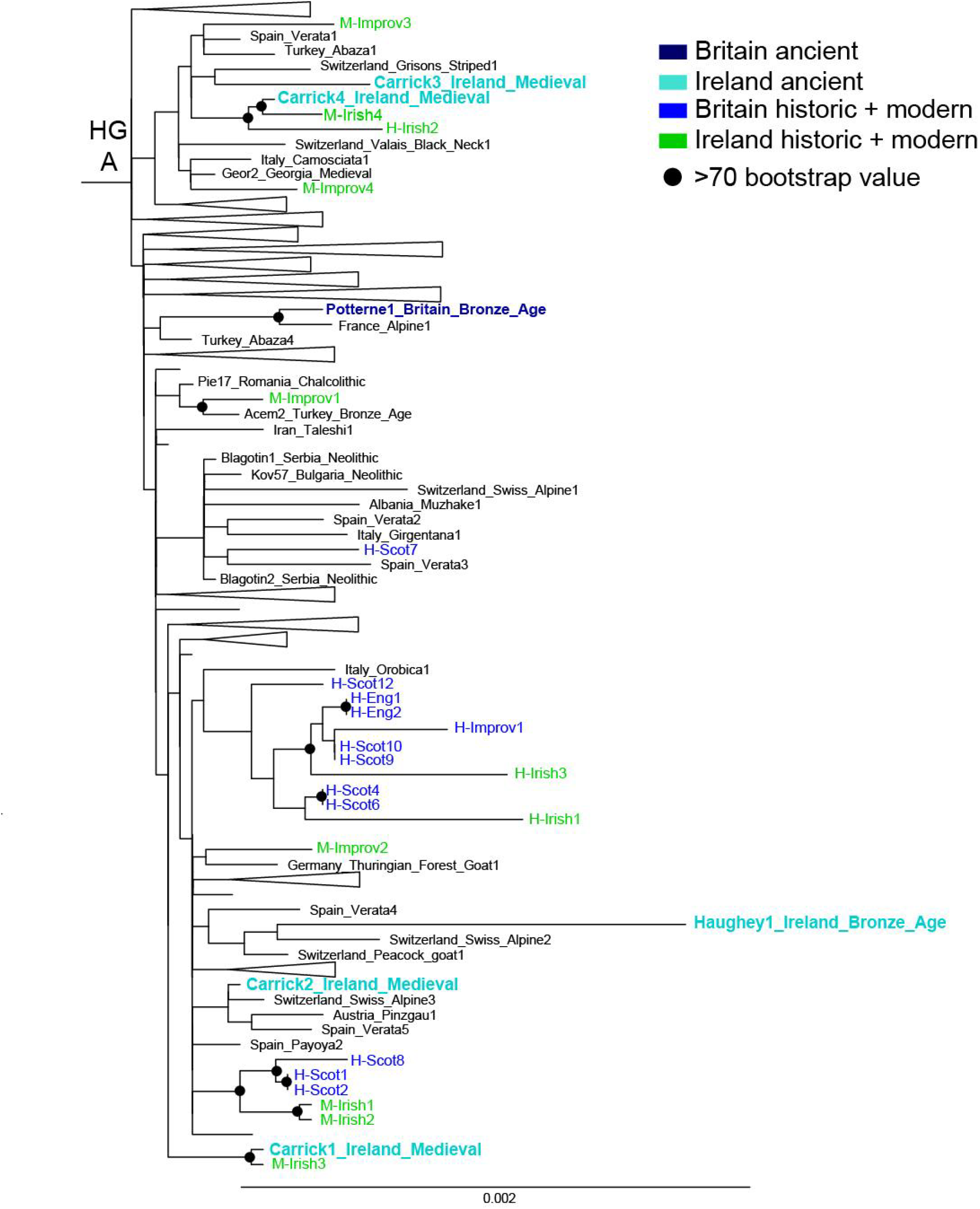
Goat mtDNA haplogroup A phylogeny, including ancient, historic, and modern sequences. Haplogroup A clade shown above was extracted from a phylogeny of all *C. hircus* and *C. aegagrus* mtDNA sequences (Figure S12). Phylogeny includes historic and modern sequences of British and Irish goats reported in (Cassidy et al., 2017).

### 3.5 Inbreeding

Finally, we assessed evidence of inbreeding and long-term population size in Irish goats by runs-of-homozygosity (ROH) analysis. Using imputed ancient genomes with sufficient coverage (>1.2 million SNPs), we found variation of inbreeding profiles between two Carrickfergus goats (Figure 5). Carrick3 shows a low cumulative sum of ROH, with only a small fraction of ROH >4Mb, thus lacking signals of recent inbreeding. In contrast, Carrick1 shows a high cumulative sum of ROH, with several long ROHs indicative of recent inbreeding. The Carrick1 ROH profile is similar to modern Old Irish Goats, (Figure 5), which all show a high cumulative sum of ROH. In particular, evidence of ROH from the shortest to longest length category (>16Mb) are suggestive of a recent history of demographic decline in Old Irish Goats, possibly exacerbated by isolation on the island (Bertolini et al., 2018; Cardoso et al., 2018; Petretto et al., 2024). However, the elevated cumulative sum of ROH >16 Mb in Carrick1 suggests relatively recent inbreeding. Assessing between-individual chromosomal sharing (identity-by-descent or IBD analysis), we find low sharing between Carrick1 and Carrick3, with only one shared segment of 7.2 cM suggesting no recent common ancestors between the two Carrickfergus samples (Table S9). We find no evidence of chromosomal segment sharing between any of the Carrick samples and a Bronze Age British goat (Potterne1).

**Figure 5.**
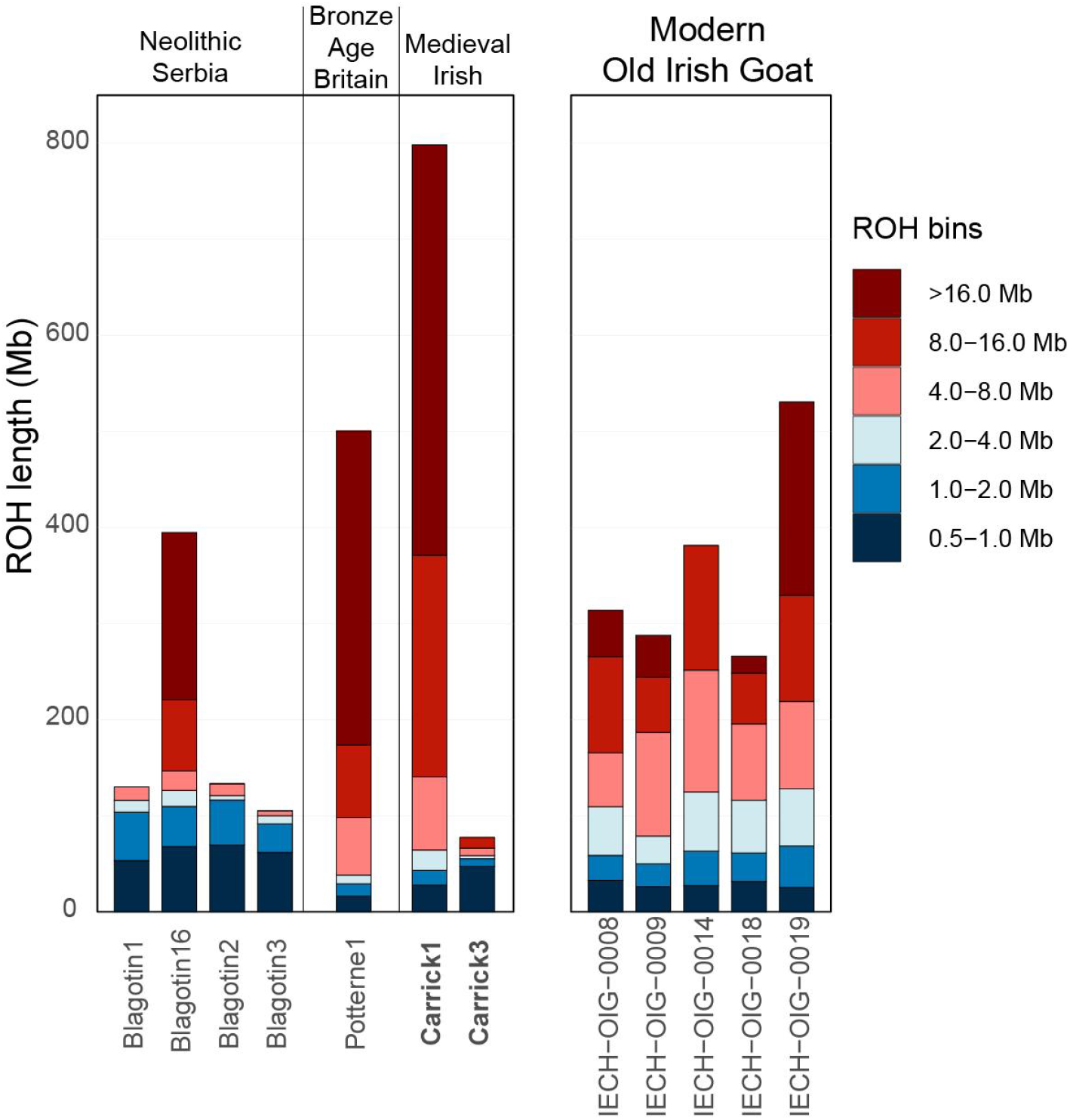
Run-of-homozygosity (ROH) analysis of ancient Irish and British goats, and the Old Irish Goat breed today. Each bar represents a genome, stratified by the total Mb amount of genome falling within a ROH/inbreeding tract of a certain length.

## 4. Discussion and Conclusion

It remains difficult to differentiate between sheep and goats using morphological and metrical approaches, but it was necessary to undertake such analyses to target individual goats for further genetic analysis. Verification of the osteological approaches is increasingly being undertaken using ZooMS to enable a greater understanding to be gained of past sheep and goat husbandry practices (Buckley et al., 2010; Seabrook et al., 2025). This approach was successful in the current study and enabled both Late Bronze Age and Late medieval goats to be identified. The four goats from Haughey’s Fort are of particular significance since they are the oldest confirmed goat remains in Ireland, with the radiocarbon dating of Haughey1 confirming unequivocally their Late Bronze Age provenance (1,112-947 cal BCE).

Importantly, assessing both proteomic and genetic evidence allowed us to confirm the species of more specimens (initially identified through archaeozoological methods as likely domestic goats or whose species was inconclusive) than either method alone. This illustrates the complementary nature of these methodologies: the relatively-inexpensive ZooMS offers inferential power for specimens in which no DNA can be recovered, while genetic analyses, when feasible, offer a rich demographic analysis.

Genomic analyses from even a small number of ancient or historic individuals can shed light on the history of more recent populations. Here, we demonstrate its utility in recontextualizing our understanding of the Old Irish Goat breed. The high genetic connectivity (Figure 3, S5) observed between the Old Irish Goats and the goats of both Late Bronze Age Haughey’s Fort and Late medieval Carrickfergus suggest substantial population continuity of goats within the island of Ireland.

### 4.1 The goats of Haughey’s Fort and Carrickfergus

The deep signal of genetic connection between the Late Bronze Age Haughey’s Fort goat and the Old Irish Goat of today (Figure 3), albeit inferred from a small number of aligned reads, is consistent with a persistence of the indigenous goat population on the island of Ireland over 3,000 years. Continuity of local goat populations has been reported elsewhere in Europe (Rannamäe et al., 2023), and more broadly in sheep (e.g. (Rannamäe et al., 2016; Rossi et al., 2021)). It is likely that in most cases, excluding where historic texts provide alternative evidence, local breeds represent distant descendants of herds introduced to that region hundreds or thousands of years previously.

As the two Carrickfergus medieval goats have notably different inbreeding profiles (Figure 5) and lack substantial numbers of shared IBD chromosomal segments, it is plausible they had different origins or derived from different incipient breeds. The former of these is possible given the suggestions of the importation of goat horn cores during this period (McCormick and Murray, 2017), and exchange of livestock animals in the Early medieval period (McCormick, 1992). However, we find no evidence of differing genetic ancestry between them, using *D* statistics (Table S8). We cautiously interpret this to reflect the varied goat keeping practices of the Carrickfergus communities through time, of a herd stock closely related to the Irish primitive goat breed today. However, it is possible that the Carrick1 individual dates to a more recent time than Carrick3, given the differences in genetic signal and read length distributions (Figure S3), leading to diachronistic patterns of genomic ancestry rather than diversity of herd genetics.

In this context, there is evidence of the long-distance trade of materials derived from goats during the Late medieval period. Hides (including goat hides) are thought to be one of the major export goods from Ireland at this time and were principally exported from Dublin, Drogheda and Carrickfergus itself (Longfield, 1929). The wills of two Coventry merchants who died in Dublin in the 15th century include mention of goat skins (O’Neill, 1987), as do the 1518 accounts of Bristol Customs, detailing goat and kid skin imports from Ireland (Longfield, 1929). As Carrickfergus was one of the most important Irish ports up until the early 17th century (McSkimin, 1909), it is quite probable that there would have been a demand for goat skins for export purposes. However, our knowledge of the herds necessary to produce such goat skins in the medieval period is limited, with little data available from rural contexts (McCormick and Murray, 2017). There are hints in the historical sources, however, that large quantities of goats were at least occasionally being kept. In 1282, the Ulster Anglo-Norman lord, FitzWarin, for example, is reported to have lost 2,000 two-year-old hogs and goats (McNeill, 1980). Given the predominance of females in the four Carrickfergus samples analyzed here, it is tempting to speculate that these were kept local as a milk source (McCormick and Murray, 2017). Indeed, Offence Number 39 of a paper presented to each Quarter Session of the Grand Jury of Carrickfergus prior to 1692 stated that it was an offence to ‘keep any cows, calves, sheep or goats either standing in the streets, church yard or at the stand within the key’ (McSkimin, 1909), suggesting that goats were commonly found within the town. It is also possible that some goats were brought from a rural locale to Carrickfergus as part of trade activities.

### 4.2 The history of Irish goat

We note that the available nuclear and mitochondrial genomes of North European goats is relatively limited. In particular, this restricts our ability to disentangle patterns of matrilineal, mitochondrial DNA. Our historic and ancient mitochondrial sequences (Figure 4) show substantial diversity within haplogroup A, and recontextualize several published historic and modern Irish goat mitochondrial sequences (Cassidy et al., 2017) as grouping within Irish-specific matrilines. However, mitochondrial sequences from Northern Europe may recontextualize these relationships, such as the potential connectivity between Irish and Scandinavian Viking goat herds (Manunza et al., 2023). Similarly, nuclear genomes from British and Irish breeds such as Bilberry, Aran, Cheviot, Golden Guernsey, Bagot, and English, Scottish, and Welsh Old/primitive goats would greatly enhance our understanding of their collective genetic history, in addition to aiding conservation efforts. Indeed, the population collapse of Old Irish Goats — from about seven thousand individuals to a few dozen — over the last three decades (“FAOSTAT,” 2018) is evident in their genomes (Figure 4, (Cardoso et al., 2018; Petretto et al., 2024)). As the same inbreeding patterns are not observed in both of the two Carrickfergus goats, we infer the genetic decline is primarily a recent phenomenon; additional pre-modern genomes would allow this demographic history to be more fully resolved.

## Acknowledgements

Sadly, Dr. Judith Findlater passed away on 18 December 2023 following a battle with cancer. We are very grateful to her mother, Philomena, and brother, Michael, for sharing her research materials with us.

Thanks are due to Damien Houlahan and Mark Nugent of the Navan Centre, Armagh City, Banbridge and Craigavon Borough Council, for help accessing the goat horncore from Haughey’s Fort and to Jackie McDowell, Historic Environment Division, Department for Communities, for permission to sample the bones from Carrickfergus. We thank Sinead Keane and Sean Caroloan for access to the Old Irish Goat image. We thank Daniel Bradley for his mentorship and the use of both laboratory and computational facilities at TCD.

The ZooMS analysis was carried out by Dr Samantha Greeves at BioArCh, University of York, who gratefully acknowledges the use of the UltrafleXtreme MALDI-ToF/ToF instrument in the York Centre of Excellence in Mass Spectrometry. The Centre was created thanks to a major capital investment through Science City York, supported by Yorkshire Forward with funds from the Northern Way Initiative, and subsequent support from EPSRC (EP/K039660/1; EP/M028127/1). V.E.M. was supported by the European Research Council under the European Union’s Horizon 2020 research and innovation program (885729-AncestralWeave). A.S. was supported by the European Research Council under the European Union’s Horizon 2020 research and innovation program (101055195-ANSOC). This publication has emanated from research conducted with the financial support of Taighde Éireann – Research Ireland under Grant numbers 21/PATH-S/9515(T) and GOIPG/2023/4390. This work was partly funded through the AHRC Northern Bridge Doctoral Training Partnership through a Collaborative PhD Studentship (Feeding Medieval Carrickfergus – A Multi-proxy Study of Livestock Husbandry in a Frontier Town) awarded to Judith Findlater.

## Data availability

The VarGoats genotypes are under embargo until 31st December 2025, and will then be available at ENA under Project accession PRJEB90141. The genetic data will be deposited in the European Nucleotide Archive (ENA; accession no. PRJEB98255). Previously published data was used for this work (Daly et al., 2021, 2018).

