## Supplementary_Note_Figures_S1-12 for "Old Goats: 3,000 years of genetic connectivity of the domestic goat in Ireland"

**Supplementary Note**
Given the low-coverage of the Late Bronze Age Irish goat (Haughey1), a downsampling approach of high-coverage ancient goats was employed to determine if this could influence outgroup $f_3$ statistics. The number of downsampled sites ranged from very low (5000, 10000, 20000), to low (40000, 80000, 100000) to medium (200000, 500000, 1000000). Outgroup $f_3$ calculated from the high-coverage individual were compared to the downsampled replicates (Table S10).

Goats with European ancestry, Blagotin3 (Neolithic Serbia) and Potterne1 (Bronze Age Britain) show a high correlation (>0.85) in outgroup $f_3$ when tested for the lowest number of sites (5000), this increases to >0.93 when increasing the sites to 10,000. This is also the case for Semnan3 (Neolithic Iran), which shows a correlation of 0.89 and 0.91 for 5000 and 10,000 sites, respectively. This highlights that outgroup $f_3$ of low-coverage ancients from a European or Asian origin are robust. In contrast, ancestries less represented in the VarGoats dataset show lower correlation for downsampled replicates. The medieval Levantine goat (Yoqneam2) shows a low correlation (<0.70) for 5000 and 10,000 sites, which increases to >0.92 when using 20,000 sites. A similar trend is observed for a medieval Georgian goat (Kazbegi1) and for a Epipaleolithic wild goat from Direkli cave (Direkli1-2). They show a correlation >0.90 for 80,000 and 40,000 sites, for Kazbegi1 and Direkli1-2 respectively. The individual with the worst scoring correlation is Acem2, a Bronze Age Turkish goat, which has a correlation of 0.12 with the lowest number of sites (5000), and only achieves a consistent correlation of >0.90 at 500,000 sites.

While calculating outgroup $f_3$ statistics on low-coverage samples is feasible, caution is warranted when the number of overlapping sites is very low, particularly if the sample's ancestry is underrepresented in the comparative dataset.

### Supplementary Figures

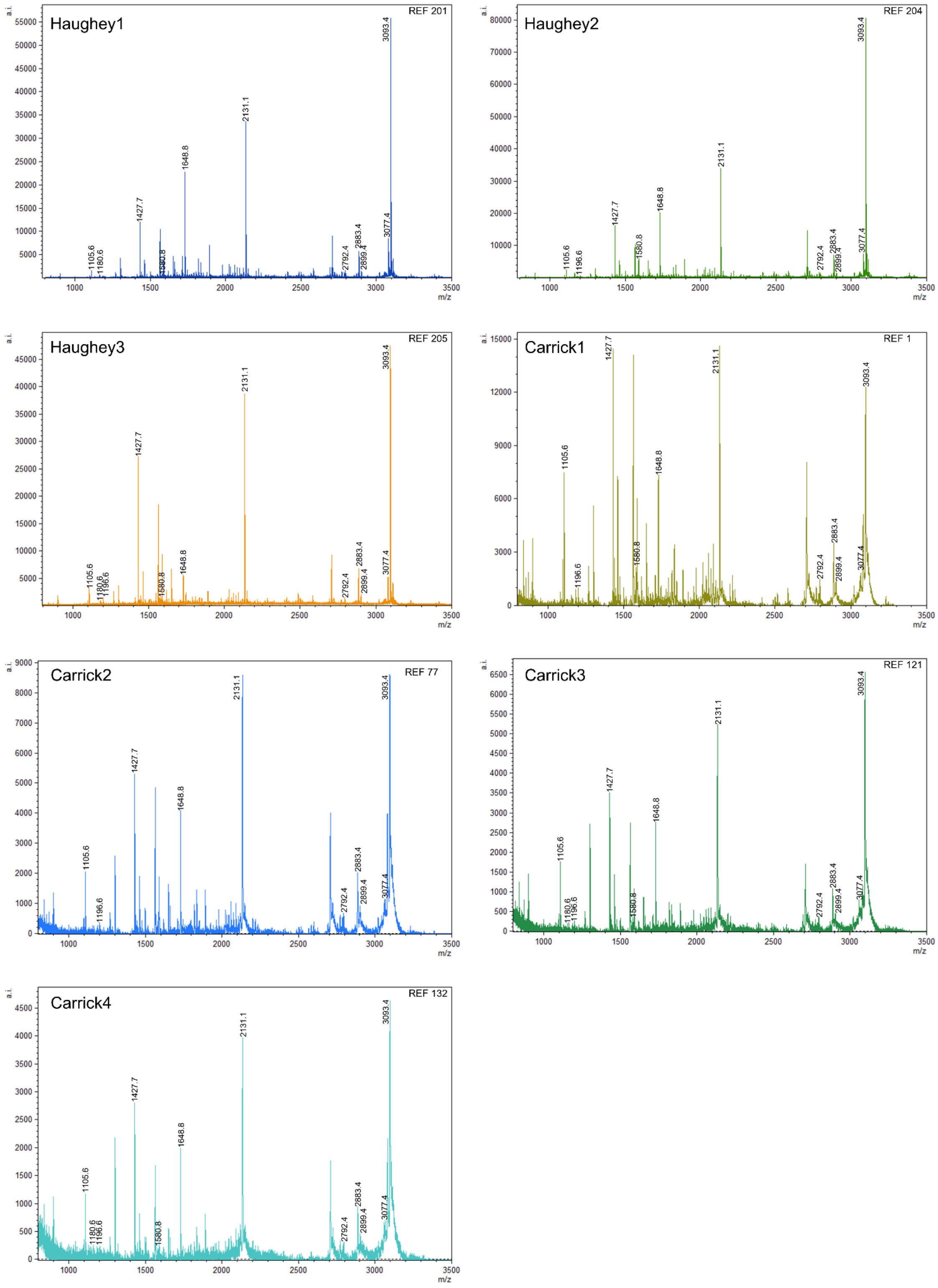


Figure S1: ZooMS peptide mass fingerprint spectra of Haughey’s Fort (Haughey[1-3]) and Carrickfergus (Carrick[1-4]), compared against reference spectra. Presence of 3093 m/z and absence of 3033 m/z are consistent with *Capra hircus*.

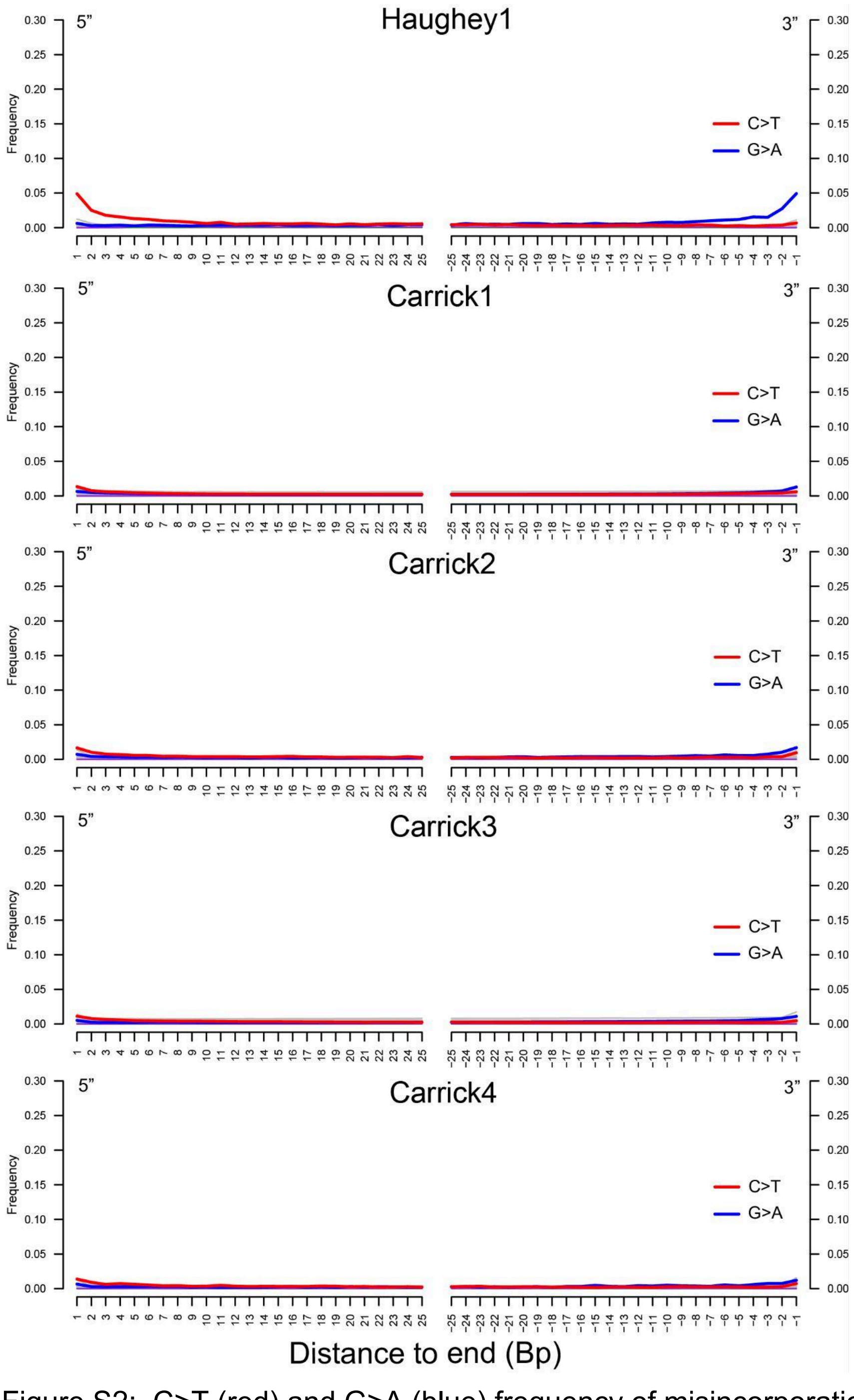


Figure S2: C>T (red) and G>A (blue) frequency of misincorporation rates of ancient Irish goat samples, at the 3’ and 5’ end of reads. All samples were Uracil DNA gylcosylase treated.

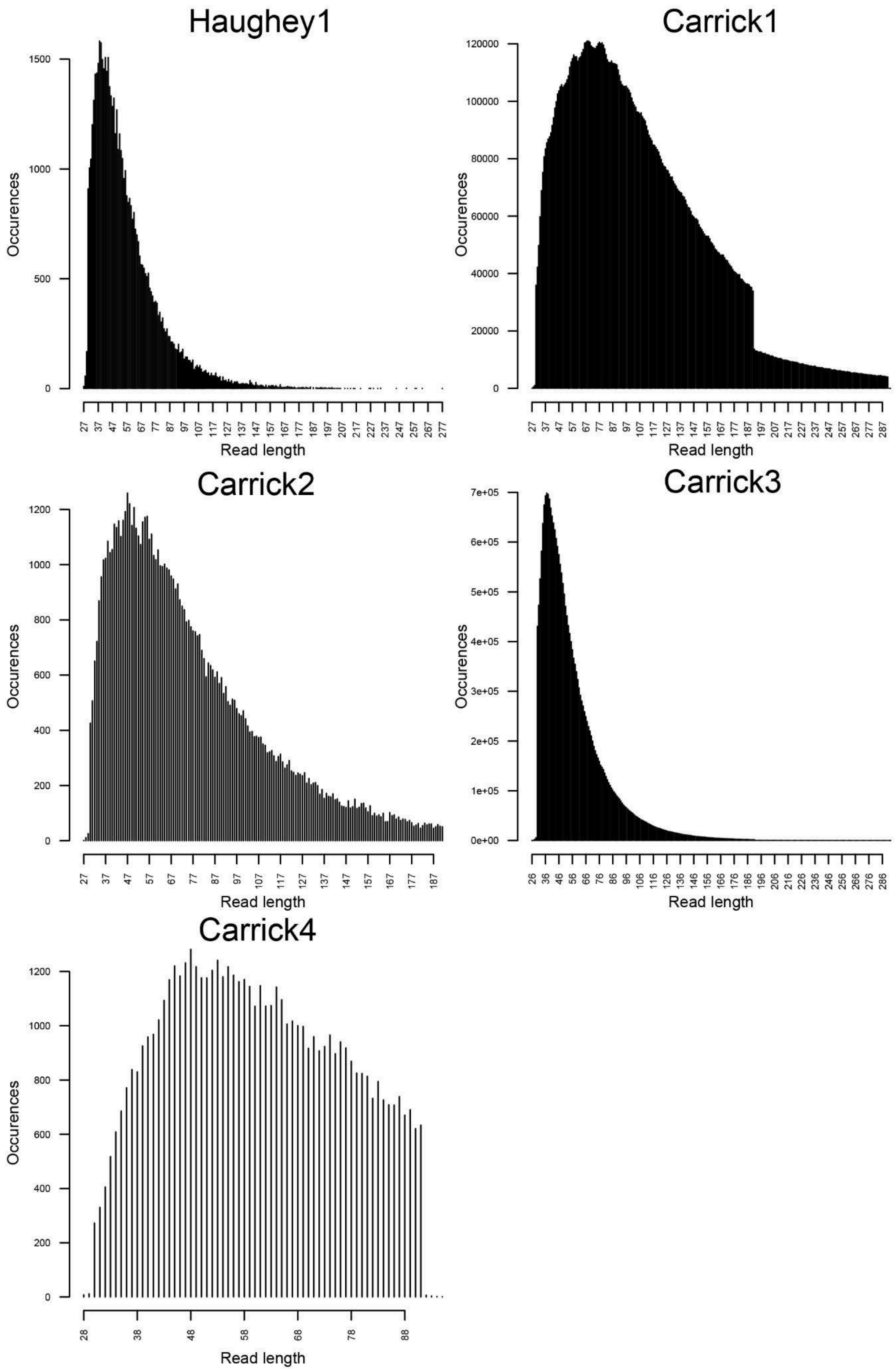


Figure S3: mapDamage read length distribution of ancient Irish goat samples.

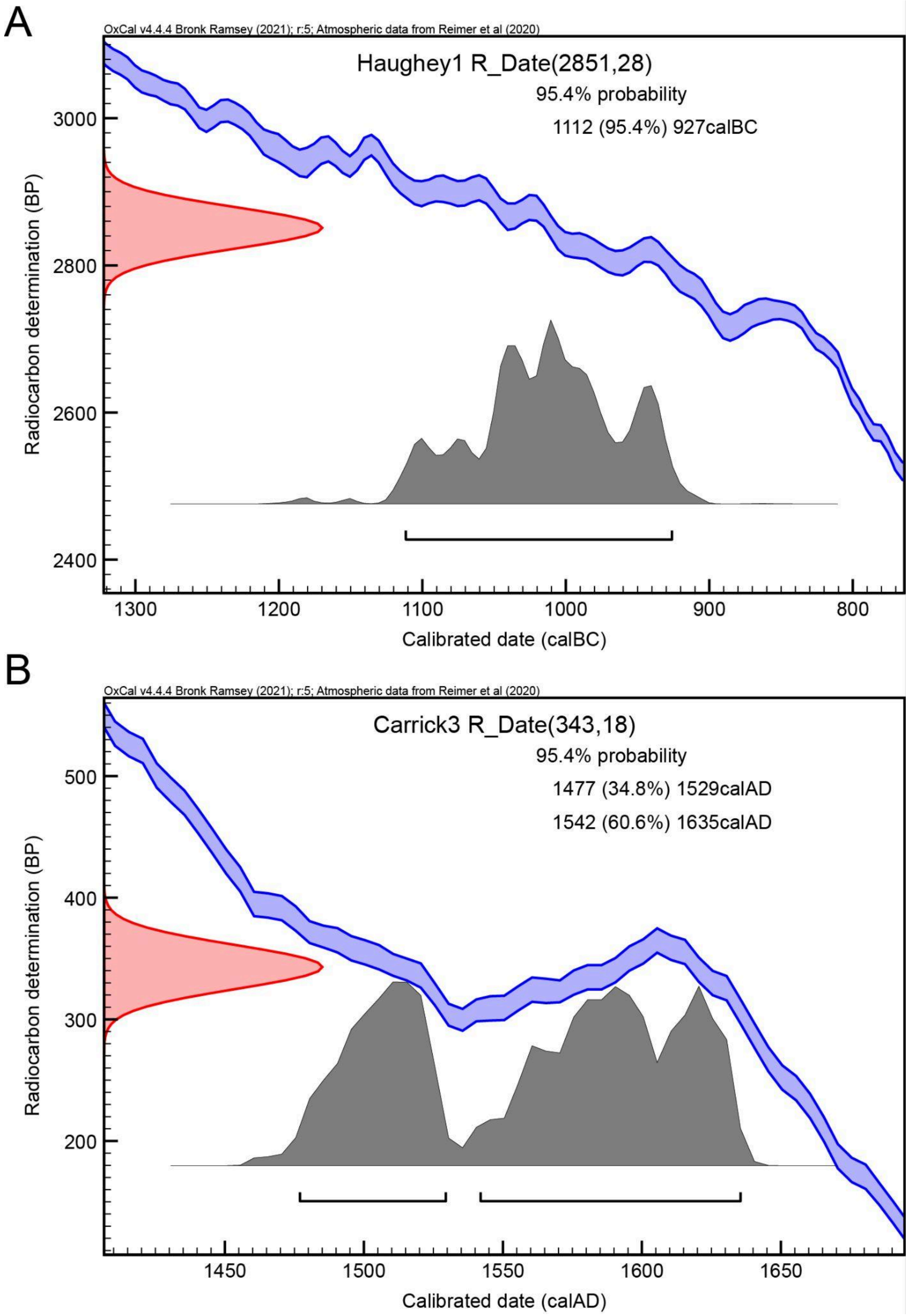


Figure S4 Radiocarbon calibration curve for A) Haughey1 and B) Carrick3.

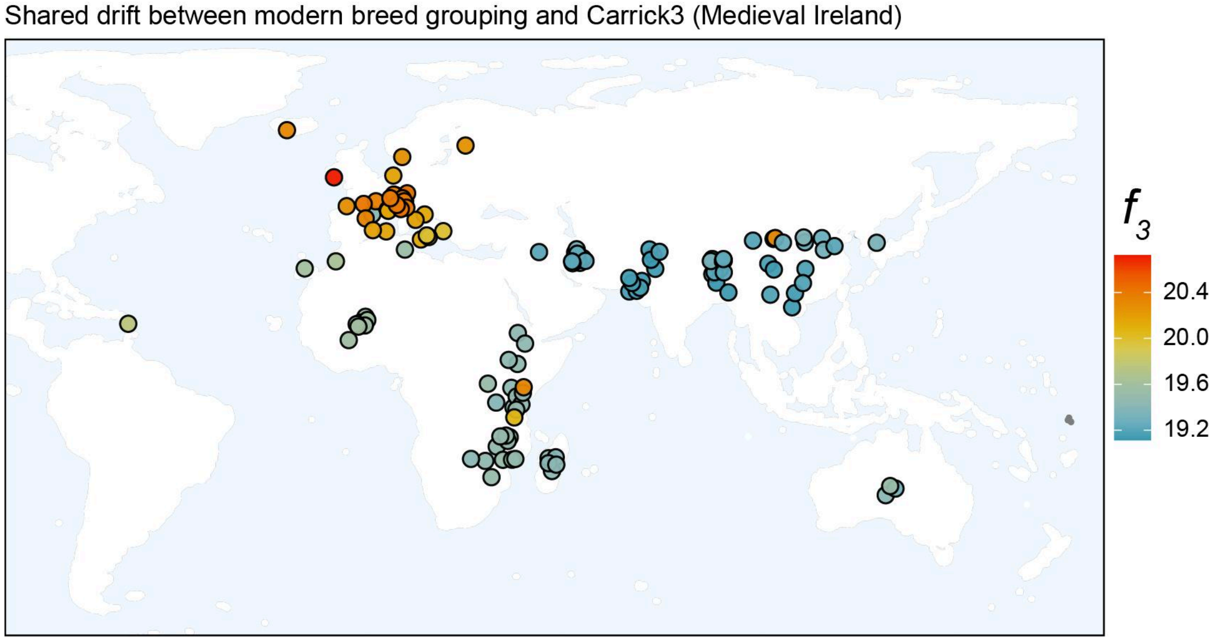


Figure S5: Geographical representation of outgroup $f_3$ values of a medieval goat from Carrickfergus, Ireland (Carrick3) and modern domestic goat breeds, measuring the relative shared drift between Carrick3 and modern breeds.

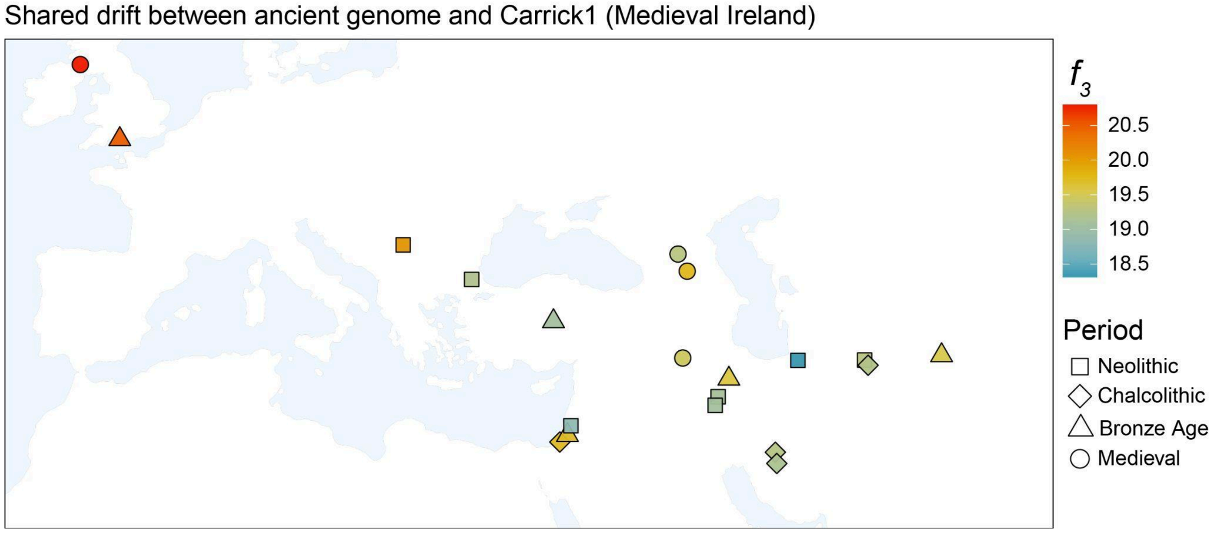


Figure S6: Geographical representation of outgroup $f_3$ values of a medieval goat from Carrickfergus, Ireland (Carrick1) and other ancient goats, measuring the relative shared drift between Carrick1 and other ancient goats.

Shared drift between ancient genome and Carrick3 (Medieval Ireland)

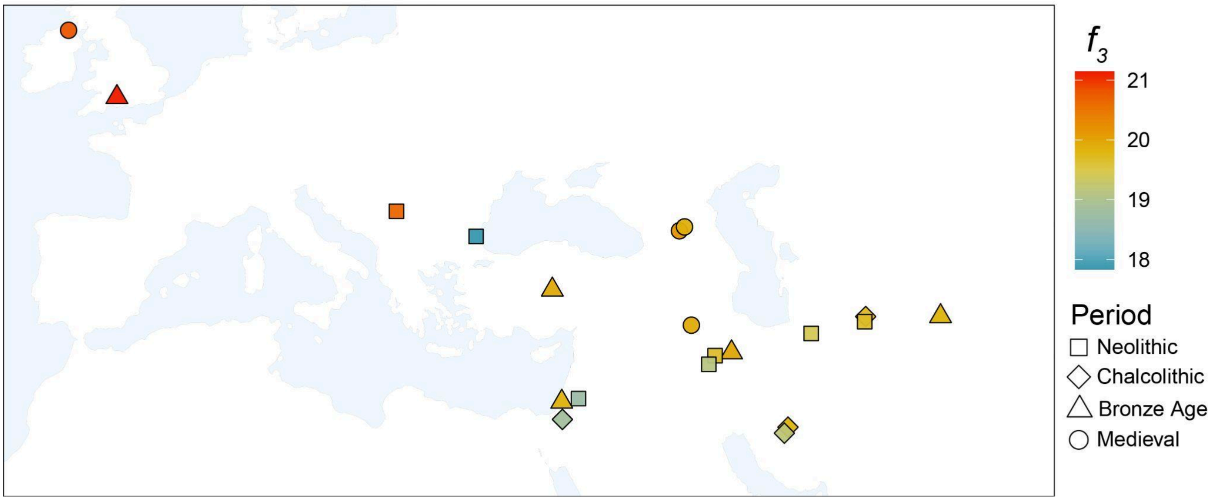


Figure S7: Geographical representation of outgroup $f_3$ values of a medieval goat from Carrickfergus, Ireland (Carrick3) and other ancient goats, measuring the relative shared drift between Carrick3 and other ancient goats.

Shared drift between ancient genome and Haughey1 (Bronze Age Ireland)

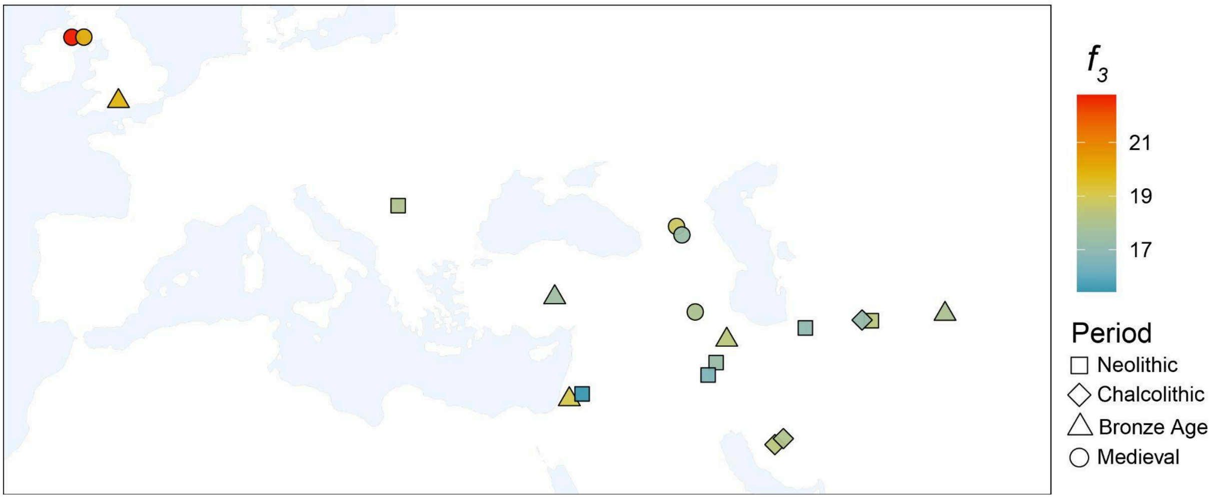


Figure S8: Geographical representation of outgroup $f_3$ values of a Bronze Age goat from Haughey's Fort, Ireland (Haughey1) and other ancient goats, measuring the relative shared drift between Haughey1 and other ancient goats.

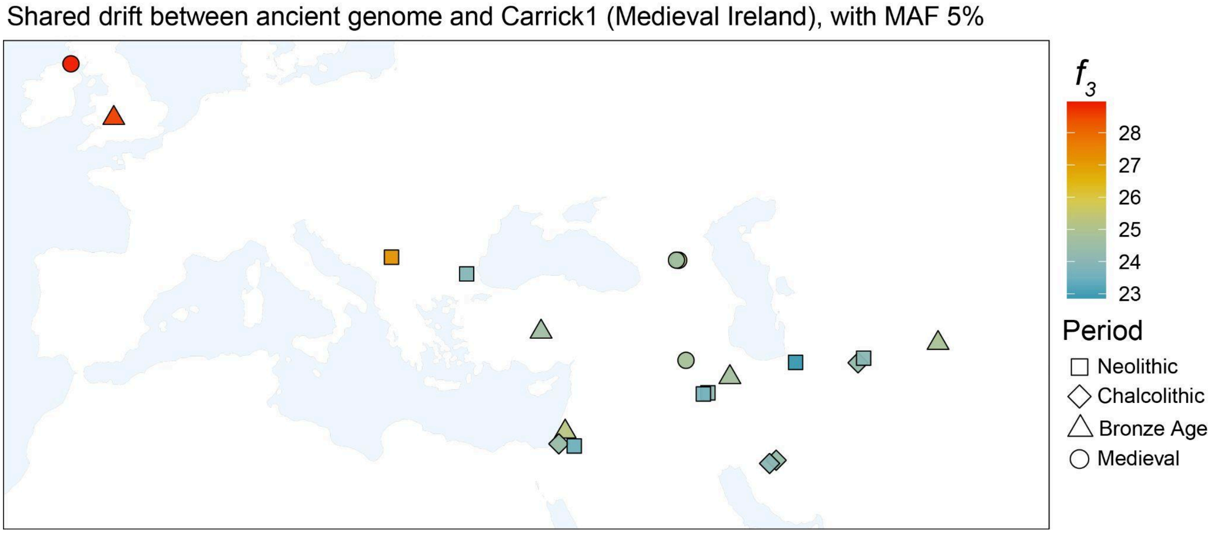


Figure S9: Geographical representation of outgroup $f_3$ values of a medieval goat from Carrickfergus, Ireland (Carrick1) and other ancient goats, measuring the relative shared drift between Carrick1 and ancient goats. Pseudohaploid genotypes were filtered for a MAF >5%.

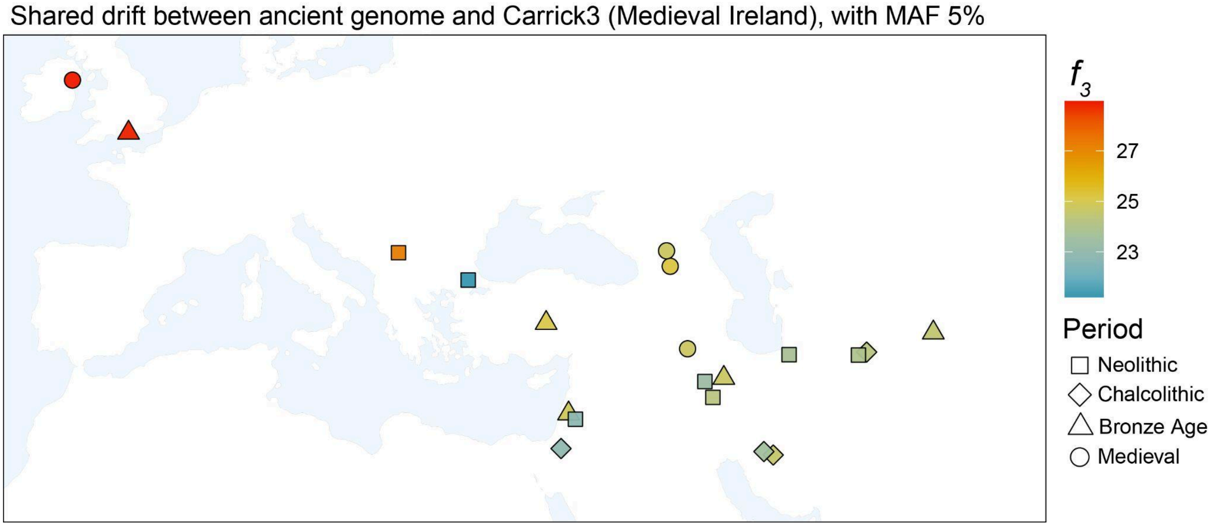


Figure S10: Geographical representation of outgroup $f_3$ values of a medieval goat from Carrickfergus, Ireland (Carrick3) and other ancient goats, measuring the relative shared drift between Carrick3 and ancient goats. Pseudohaploid genotypes were filtered for a MAF >5%.

Shared drift between ancient genome and Haughey1 (Bronze Age Ireland), with MAF 5%

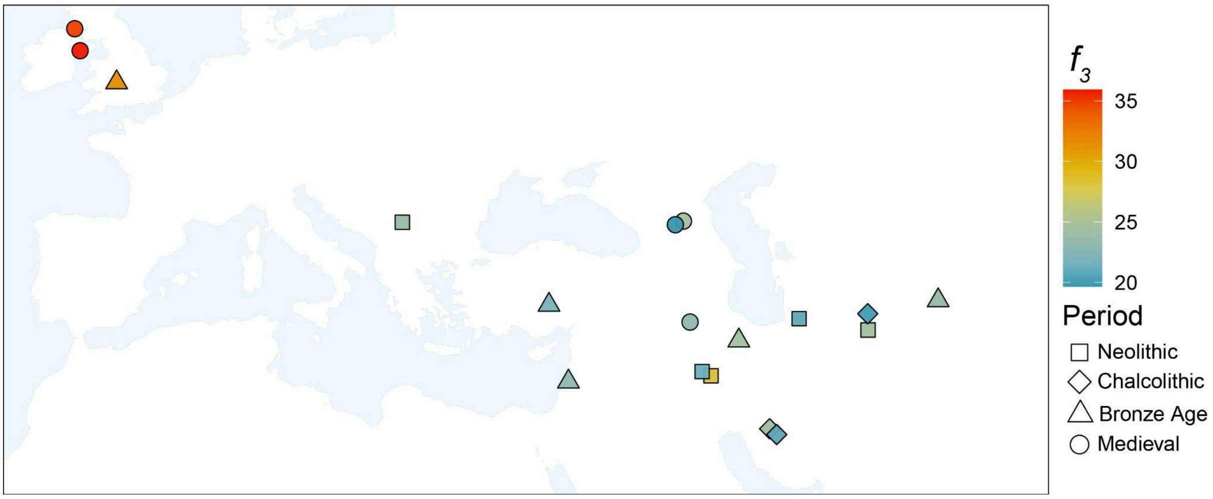


Figure S11: Geographical representation of outgroup $f_3$ values of a Bronze Age goat from Haughey's Fort, Ireland (Haughey1) and other ancient goats, measuring the relative shared drift between Haughey1 and ancient goats. Pseudohaploid genotypes were filtered for a MAF >5%.

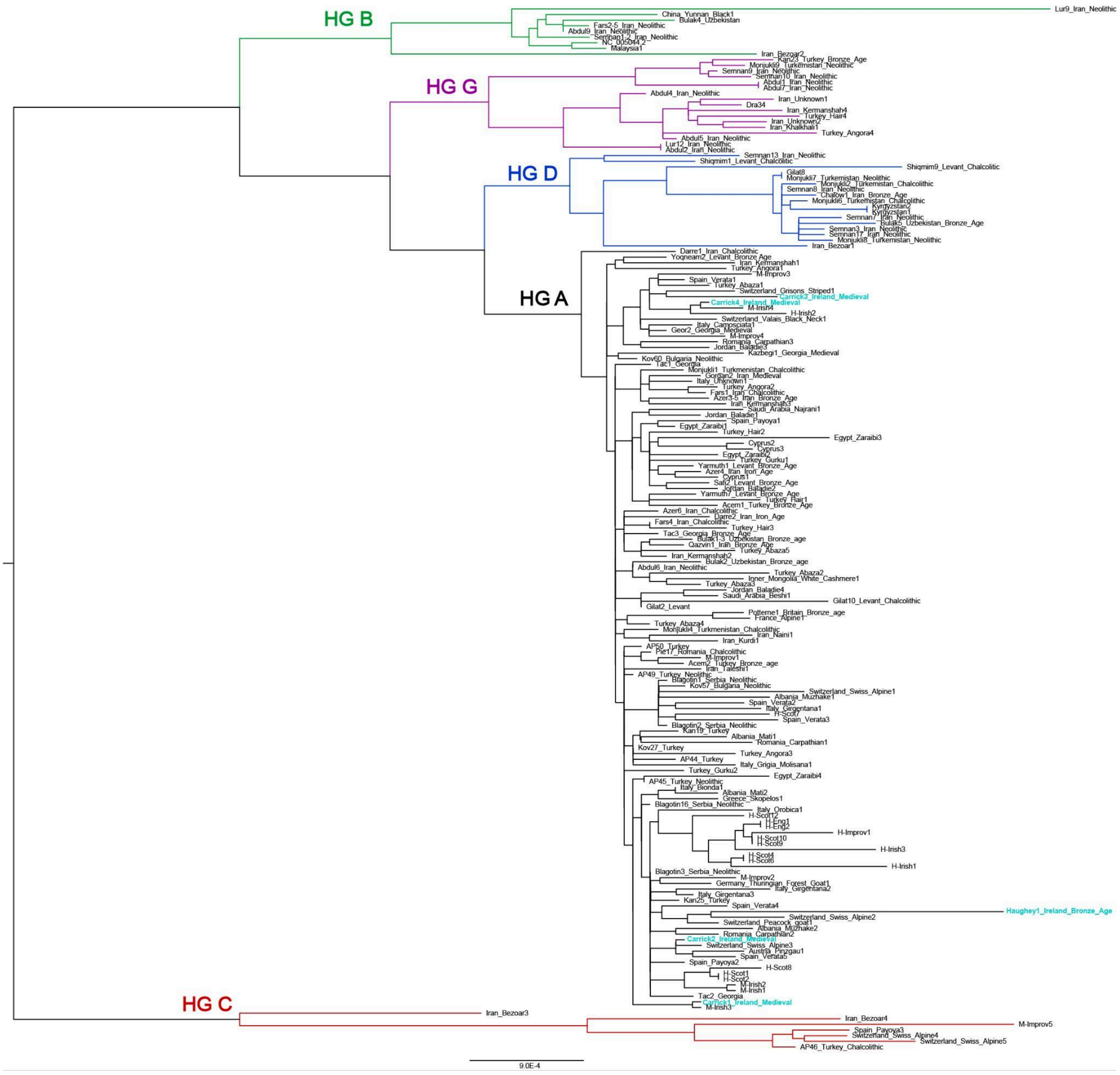


Figure S12: *C. hircus* and *C. aegagrus* mtDNA phylogeny, including ancient, historic, and modern sequences. Samples introduced in this study are highlighted in light blue.
